# Cofactor selectivity in methylmalonyl-CoA mutase, a model cobamide-dependent enzyme

**DOI:** 10.1101/637140

**Authors:** Olga M. Sokolovskaya, Kenny C. Mok, Jong Duk Park, Jennifer L. A. Tran, Kathryn A. Quanstrom, Michiko E. Taga

## Abstract

Cobamides, a uniquely diverse family of enzyme cofactors related to vitamin B_12_, are produced exclusively by bacteria and archaea but used in all domains of life. While it is widely accepted that cobamide-dependent organisms require specific cobamides for their metabolism, the biochemical mechanisms that make cobamides functionally distinct are largely unknown. Here, we examine the effects of cobamide structural variation on a model cobamide-dependent enzyme, methylmalonyl-CoA mutase (MCM). The *in vitro* binding affinity of MCM for cobamides can be dramatically influenced by small changes in the structure of the lower ligand of the cobamide, and binding selectivity differs between bacterial orthologs of MCM. In contrast, variations in the lower ligand have minor effects on MCM catalysis. Bacterial growth assays demonstrate that cobamide requirements of MCM *in vitro* largely correlate with *in vivo* cobamide dependence. This result underscores the importance of enzyme selectivity in the cobamide-dependent physiology of bacteria.

## Introduction

Cobalamin, commonly referred to as vitamin B_12_, is a versatile enzyme cofactor used by organisms in all domains of life. In humans, cobalamin is essential for methionine synthesis and the breakdown of fatty acids, amino acids, and cholesterol [1, 2]. Bacteria and archaea additionally use cobalamin and related cofactors, cobamides, for deoxyribonucleotide synthesis [3], metabolism of various carbon and energy sources [4-17], synthesis of secondary metabolites [18-25], sensing light [26], and other processes [15-17, 27-32]. The finding that 86% of bacterial species encode at least one cobamide-dependent enzyme in their genome [33] demonstrates the prevalence of cobamide-dependent metabolisms. Widespread use of these cofactors can be attributed to their chemical versatility, as they facilitate challenging chemical reactions including radical-initiated rearrangements, methylation reactions, and reductive cleavage of chemical bonds [34, 35].

All cobamides share the same core structure (Figure 1): a corrin ring that coordinates a cobalt ion, a variable “upper” axial ligand (*R* in Figure 1), and a pseudo-nucleotide that is covalently attached to the corrin ring through an aminopropanol linker [36] or an ethanolamine linker, in the case of nor-cobamides [37, 38]. The major differences among cobamides are in the structure of the nucleotide base, more commonly referred to as the lower axial ligand for its ability to coordinate the central cobalt ion. In cobalamin, the lower ligand is 5,6-dimethylbenzimidazole (Figure 1, boxed); in other cobamides, different benzimidazoles, phenolics, and purines constitute the lower ligand (Figure 2C) [39-43].

**Figure 1:**
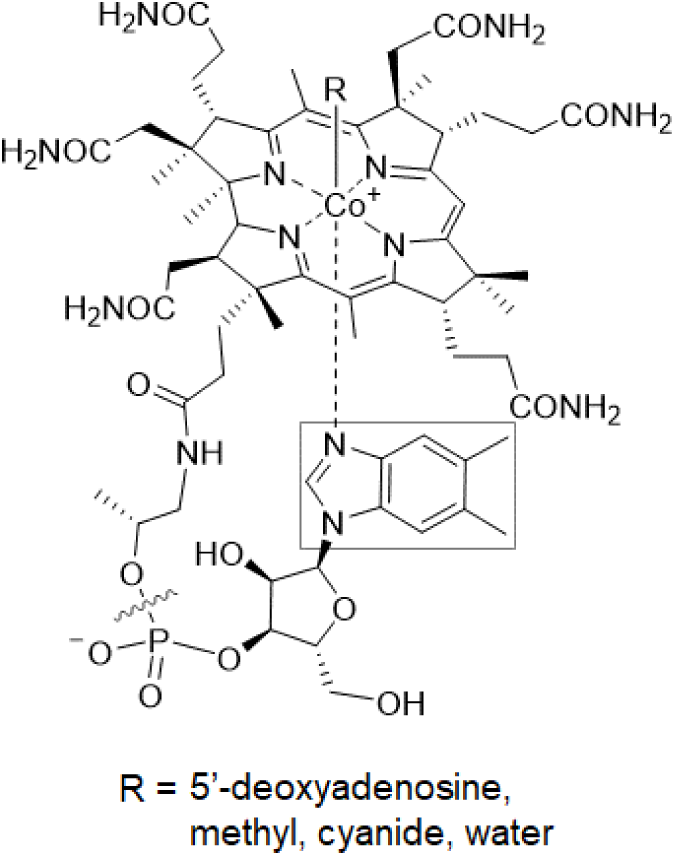
The structure of cobalamin. The lower ligand, boxed, varies in other cobamides. Cobinamide, a cobamide precursor, lacks a nucleotide base (delineated by the wavy line).

**Figure 2:**
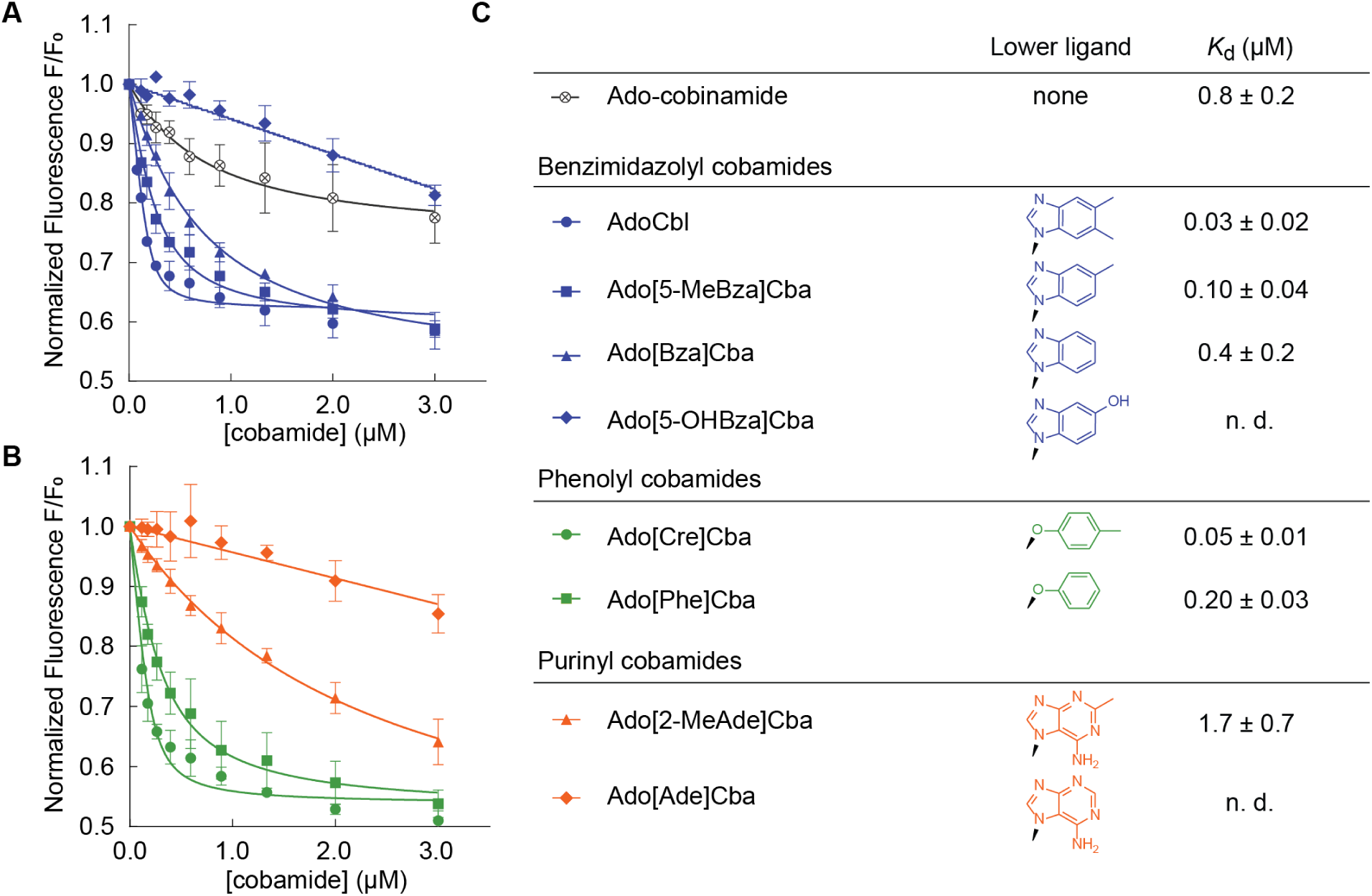
Binding of diverse cobamides to *Sm*MCM (see also Figure S1). Fluorescence decrease of *Sm*MCM when reconstituted with (A) benzimidazolyl cobamides (blue) and cobinamide (black), and (B) phenolyl (green) and purinyl (orange) cobamides. Data points represent the mean and standard deviation of three technical replicates from a single experiment. (C) *K*_d_ values for different cobamides, reported as the average and standard deviation of three or more independent experiments, each consisting of technical triplicates. “n. d.,” not determined, indicates that binding was too weak to determine *K*_d_.

While cobamides containing different lower ligands share the same chemically reactive moieties, specifically the cobalt center and methyl or 5’-deoxyadenosyl upper axial ligands, they are nonetheless functionally distinct. Culture-based studies have shown that only a subset of cobamides supports a given bacterial metabolism, and uptake or production of other cobamides can inhibit growth [39, 44-49]. The requirements of bacteria for particular cobamides is notable given the diversity of cobamides present in host-associated and environmental samples [40-42], coupled with the absence of *de novo* cobamide biosynthesis in more than half of bacteria [33]. Despite the biological relevance of cobamide structure, and the prevalence of cobamide use among bacteria [33, 50-52], little is understood about the biochemical mechanisms by which cobamides differentially impact microbial physiology.

The effect of lower ligand structure on the biochemistry of cobamide-dependent enzymes has been studied to a limited extent. In “base-on” enzymes, the lower ligand base coordinates the central cobalt ion of the cobamide, as drawn in Figure 1 [53-55]. Because the lower ligand is part of the catalytic center of the enzyme, lower ligand structure can influence catalysis through a variety of mechanisms [56-58], and cobamides unable to form an intramolecular coordinate bond are catalytically inactive in base-on enzymes [59, 60]. In contrast, in “base-off” enzymes the lower ligand is bound by the enzyme more than 10 Å away from the active site [20, 61-70]. In a subset of base-off enzymes, referred to as “base-off/His-on,” a histidine residue from the protein coordinates the cobalt ion in place of the lower ligand [61, 63]. Despite its distance from the reactive center, lower ligand structure affects the activity of base-off enzymes, as evidenced by the cobamide cofactor selectivity of methionine synthase [71], methylmalonyl-CoA mutase (MCM) [60, 72], reductive dehalogenases [49], and other enzymes [59, 72, 73]. However, the mechanisms by which lower ligand structure affects the biochemistry of base-off cobamide-dependent enzymes remain unclear.

As MCM is one of the most abundant cobamide-dependent enzymes in bacterial genomes [33], and one of the two cobamide-dependent enzymes in humans, we have chosen to study the cobamide selectivity of MCM as a model for base-off/His-on enzymes, all of which share a structurally conserved B_12_-binding domain [63, 74]. MCM catalyzes the interconversion of *(R)-*methylmalonyl-CoA and succinyl-CoA, a bidirectional reaction used in propionate metabolism [12, 75, 76], catabolism of branched amino acids and odd-chain fatty acids [76, 77], polyhydroxybutyrate degradation [78], secondary metabolite biosynthesis [79], and autotrophic carbon dioxide fixation [4, 80]. MCM-dependent pathways have been harnessed industrially for the bioproduction of propionate, bioplastics, biofuels, and antibiotics [81-88].

The presence of a cobamide lower ligand is required for MCM activity, as evidenced by the observation that adenosylcobinamide, a cobamide intermediate lacking a lower ligand (Figure 1), does not support MCM activity *in vitro* [89]. Three studies provide evidence that MCM is selective for cobamides with particular lower ligands. First, MCM from *Propionibacterium shermanii* was found to have different apparent K_M_ values for cobamides, increasing from AdoCbl to Ado[Bza]Cba to Ado[Ade]Cba (refer to Table 1 for full names of cobamides), and MCM from sheep had a higher apparent K_M_ for Ado[Bza]Cba than AdoCbl [72]. Second, *P. shermanii* MCM had a lower apparent K_M_ for Ado[Cre]Cba than AdoCbl [60]. Third, in *Sinorhizobium meliloti* bacteroids MCM activity was highest with AdoCbl, intermediate with Ado[Bza]Cba, and absent with Ado[Ade]Cba [90]. Each of these studies includes only one or two cobamides other than cobalamin, and understandably so; cobamides are difficult to obtain in high quantities and must be purified from large volumes of bacterial cultures. Because of this, the response of MCM orthologs to the full diversity of cobamides has not been explored, and the mechanistic basis of cobamide selectivity remains unclear.

**Table 1:**
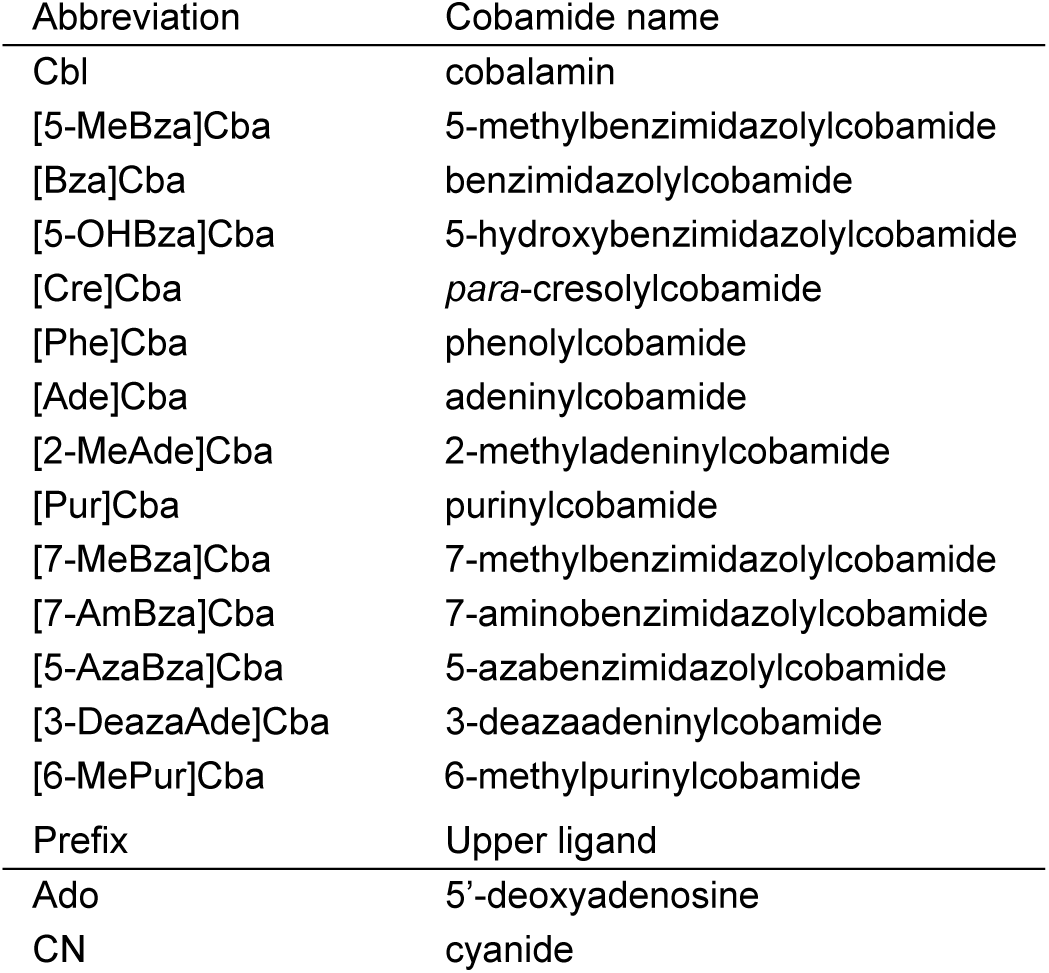
Abbreviations used for cobamides and upper axial ligands.

| Abbreviation | Cobamide name |
| --- | --- |
| Cbl | cobalamin |
| [5-MeBza]Cba | 5-methylbenzimidazolylcobamide |
| [Bza]Cba | benzimidazolylcobamide |
| [5-OHBza]Cba | 5-hydroxybenzimidazolylcobamide |
| [Cre]Cba | *para*-cresolylcobamide |
| [Phe]Cba | phenolylcobamide |
| [Ade]Cba | adeninylcobamide |
| [2-MeAde]Cba | 2-methyladeninylcobamide |
| [Pur]Cba | purinylcobamide |
| [7-MeBza]Cba | 7-methylbenzimidazolylcobamide |
| [7-AmBza]Cba | 7-aminobenzimidazolylcobamide |
| [5-AzaBza]Cba | 5-azabenzimidazolylcobamide |
| [3-DeazaAde]Cba | 3-deazaadeninylcobamide |
| [6-MePur]Cba | 6-methylpurinylcobamide |
| Prefix | Upper ligand |
| Ado | 5'-deoxyadenosine |
| CN | cyanide |

To investigate the mechanisms by which diverse lower ligands affect MCM function, we conducted *in vitro* binding and activity assays with MCM from *S. meliloti* (*Sm*MCM). We discovered major differences in the binding affinities of eight naturally occurring cobamides for *Sm*MCM, while cobamide structure affected enzyme activity to a lesser extent. Using six additional cobamides, five of which are novel analogs that have not been observed in nature or described previously, we identified structural elements of lower ligands that are determinants of binding to *Sm*MCM. To probe the hypothesis that enzyme selectivity influences bacterial growth, we characterized the cobamide dependence of *S. meliloti* growth *in vivo*. By bridging the results of *in vitro* biochemistry of three bacterial MCM orthologs and the cobamide-dependent growth phenotypes of *S. meliloti*, we have elucidated molecular factors that contribute to the cobamide-dependent physiology of bacteria.

## Results

### Lower ligand structure influences cobamide binding to MCM

We chose *Sm*MCM as a model to examine how lower ligand structure influences MCM function based on previous work demonstrating its activity as a homodimer encoded by a single gene [91, 92]. We purified eight naturally occurring cobamides for *in vitro* studies of this protein, and chemically adenosylated each cobamide to produce the biologically active form used by MCM for catalysis. Previous studies showed that binding of cobamides to *P. shermanii* MCM can be detected *in vitro* by measuring quenching of intrinsic protein fluorescence [89]. We found that the fluorescence of purified, His-tagged *Sm*MCM also decreased in a dose-dependent manner when the protein was reconstituted with increasing concentrations of AdoCbl (Figure 2A). The equilibrium dissociation constant (*K*_d_) derived from these measurements, 0.03 ± 0.02 µM (Figure 2C), is 6-fold lower than the *K*_d_ reported for *P. shermanii* MCM [89]. Ado-cobinamide also bound *Sm*MCM, as was observed with *P. shermanii* MCM [89], albeit with over 10-fold reduced affinity compared to cobalamin (Figure 2A, C).

We next measured binding of other benzimidazolyl cobamides to *Sm*MCM and found that Ado[5-MeBza]Cba and Ado[Bza]Cba, the cobamides most structurally similar to AdoCbl, also bound the enzyme. However, the absence of one or two methyl groups, respectively, in the lower ligands of these cobamides caused a decrease in binding affinity relative to AdoCbl (Figure 2A, C). Strikingly, no binding of Ado[5-OHBza]Cba to *Sm*MCM was detected at low micromolar concentrations. To rule out the possibility that Ado[5-OHBza]Cba binds *Sm*MCM but does not cause a fluorescence quench, we used an alternative, filtration-based, binding assay and observed little to no binding of Ado[5-OHBza]Cba to *Sm*MCM at micromolar concentrations (Figure S1A, B).

We expanded our analysis of *Sm*MCM-cobamide binding selectivity to include cobamides from other structural classes. Both of the phenolyl cobamides tested, Ado[Cre]Cba and Ado[Phe]Cba, bound *Sm*MCM with affinities similar to those of cobalamin and other benzimidazolyl cobamides (Figure 2B, C). In contrast, the purinyl cobamides Ado[2-MeAde]Cba and Ado[Ade]Cba had lower affinities for *Sm*MCM compared to most benzimidazolyl cobamides (Figure 2B, C): Ado[2-MeAde]Cba bound *Sm*MCM with ∼20-fold lower affinity than cobalamin, and Ado[Ade]Cba did not bind to any significant extent at micromolar concentrations (verified by the filtration assay, Figure S1C). Interestingly, for all three classes of lower ligands, the presence of a methyl substituent promoted binding relative to other cobamides of the same structural class.

### Bacterial MCM orthologs have distinct selectivity

To test whether cofactor-binding selectivity is a general phenomenon across bacterial MCM orthologs, we compared the cobamide-binding profile of *Sm*MCM to that of MCM orthologs from *Escherichia coli* (*Ec*MCM) and *Veillonella parvula* (*Vp*MCM). Activity of *Ec*MCM with AdoCbl has been reported both *in vivo* and *in vitro*, although its physiological role in *E. coli* remains unclear [82, 93]. Annotations for two MCM homologs are present in the genome of *V. parvula*, and we purified the one that exhibits MCM activity when expressed in *S. meliloti* (see Materials and Methods). Because *S. meliloti* produces cobalamin [94], *E. coli* produces [2-MeAde]Cba when provided cobinamide [95], and *V. parvula* produces [Cre]Cba [96], we expected that each ortholog should have distinct cobamide selectivity. Indeed, *Ec*MCM had highest affinity for its native cobamide, Ado[2-MeAde]Cba (Figure 3A, C). All other cobamides bound with 2- to 3-fold reduced affinities relative to Ado[2-MeAde]Cba. Similarly, *Vp*MCM had a higher affinity for Ado[Cre]Cba, its native cobamide, than AdoCbl (Figure 3B, C). *Vp*MCM also bound Ado[2-MeAde]Cba and Ado[Bza]Cba with similar affinity.

**Figure 3:**
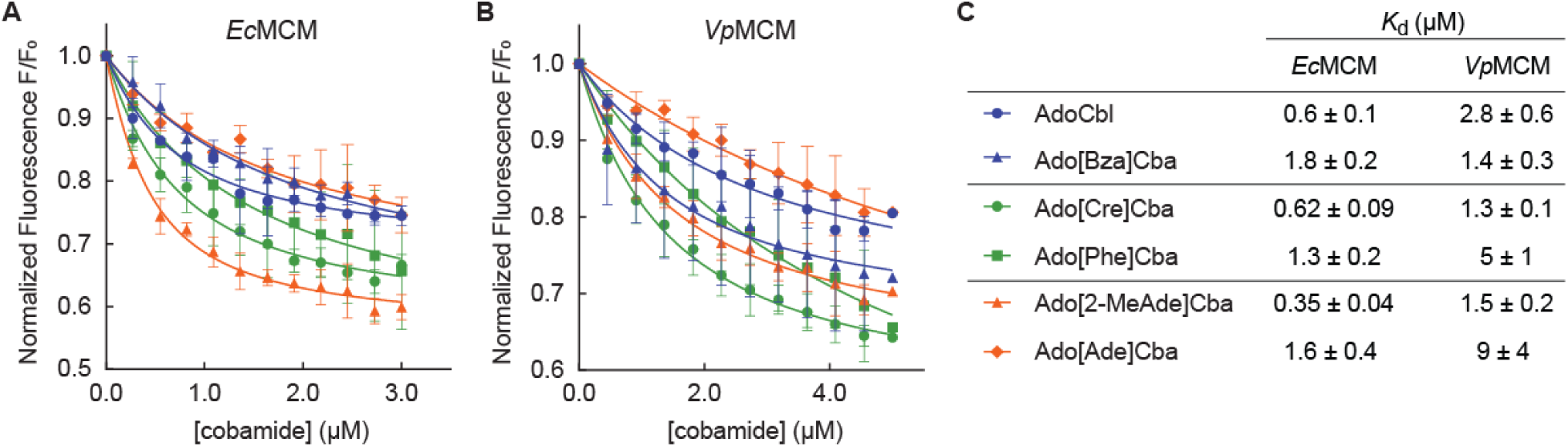
Binding selectivity of diverse MCM orthologs (see also Figure S2). Fluorescence binding assays with (A) *E. coli* MCM and (B) *V. parvula* MCM. Data points and error bars represent the mean and standard deviation, respectively, of technical triplicates from a single experiment; each replicate consisted of an independent cobamide dilution. *K*_d_ values from the fitted curves in (A) and (B) are reported in (C); error values reflect the standard error of the curve fit. *K*_d_ values for *Vp*MCM binding to Ado[Cre]Cba and AdoCbl and for *Ec*MCM binding to all cobamides were reproduced in independent experiments.

We constructed a sequence alignment of MCM orthologs from diverse organisms known to produce or use various cobamides, in search of amino acid residues that could account for differences in cobamide binding (Figure S2A). The B_12_-binding domains of diverse MCM orthologs had high overall amino acid identity (38-70%). Cases of low identity correlated with differences in the structural configuration of MCM, which occurs in different organisms as a homodimer [92, 93, 97, 98], heterodimer [61, 99-101], or heterotetramer [80, 102] (Figure S2B). We focused our analysis on residues immediately surrounding the lower ligand in the available crystal structure of *Homo sapiens* MCM [97] (*Hs*MCM) (Figure S2A, triangles). For the most part, these residues are highly conserved between orthologs. Interestingly, however, *Hs*MCM residues Phe638, Phe722, and Ala731, which are conserved in *Sm*MCM, are substituted with the more polar residues Tyr, Tyr, and Ser, respectively, in *Ec*MCM (Figure S2A), which has a higher affinity for purinyl cobamides. Introducing mutations in *Sm*MCM and *Ec*MCM to test the importance of these residues proved challenging, as it resulted in reduced protein solubility and overall impaired cobamide binding (data not shown). Whether or not these residues co-vary with cobamide selectivity across other MCM orthologs is difficult to interpret because the cobamide selectivity of MCM from most organisms is unknown.

### The lower ligand of cobamides modulates MCM reaction kinetics

We reconstituted *Sm*MCM with saturating amounts of each of the four cobamides that bound with highest affinity and measured conversion of (*R*)-methylmalonyl-CoA to succinyl-CoA under steady state conditions. Interestingly, the substrate K_M_ was nearly invariable among the cobamides tested (Figure 4). Turnover was highest with AdoCbl (26 ± 1 s^−1^) and 2- to 3-fold lower with other cobamides. Thus, all of the cobamides tested supported *Sm*MCM catalysis with modest differences in *k*_cat_. This finding is consistent with a previous observation that adenosylcobinamide-GDP, a cobamide precursor with an extended nucleotide loop and a guanine base, supported activity of *P. shermanii* MCM with only slight catalytic impairment compared to AdoCbl [103].

**Figure 4:**
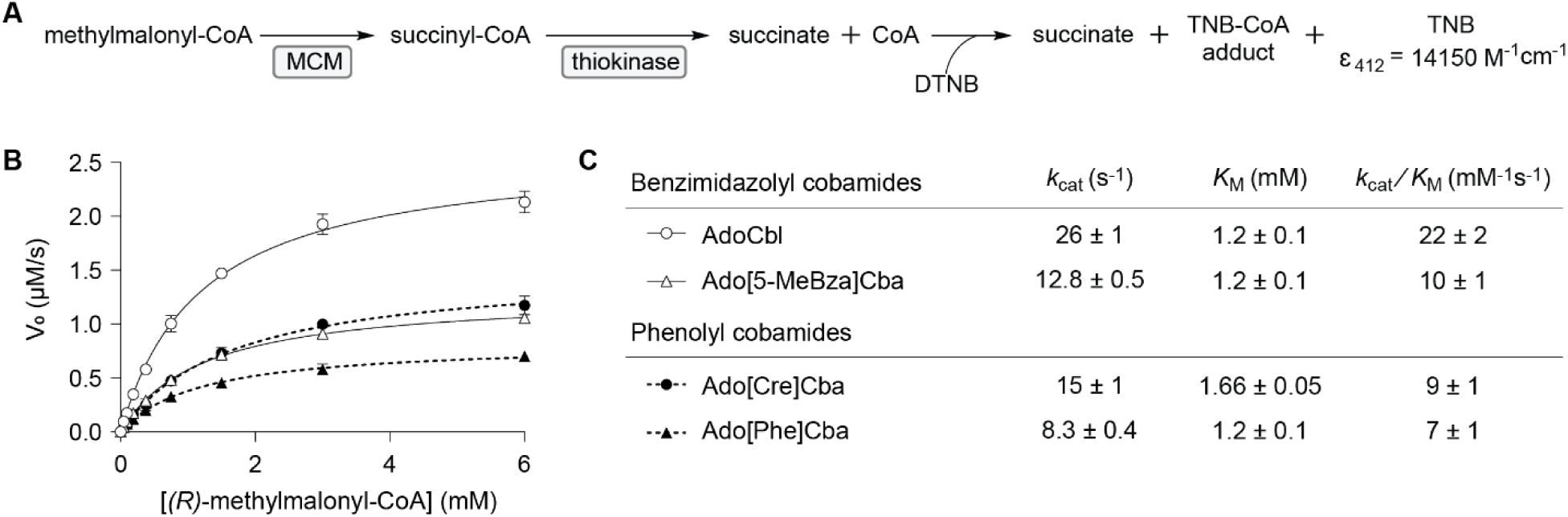
Activity of *Sinorhizobium meliloti* MCM with different cobamide cofactors. (A) Succinyl-CoA formation was detected using a coupled spectrophotometric assay [136]. DTNB, dithionitrobenzoate (Ellman’s Reagent); TNB, thionitrobenzoate; CoA, Coenzyme A. (B) Michaelis-Menten kinetic analysis of *Sm*MCM reconstituted with various cobamides. Data points and error bars represent the mean and standard deviation, respectively, of three technical replicates from one experiment; each replicate consisted of an independent substrate dilution. Kinetic constants are presented in (C).

### MCM-dependent growth of S. meliloti correlates with the binding selectivity of SmMCM for benzimidazolyl and purinyl cobamides, but not phenolyl cobamides

To assess whether the cobamide-dependent growth of *S. meliloti* reflects MCM selectivity as observed *in vitro*, we cultured *S. meliloti* under conditions that require MCM activity. Examination of metabolic pathways encoded in the *S. meliloti* genome using the KEGG database [104] suggests that the degradation of branched amino acids isoleucine and valine to succinyl-CoA, an intermediate of the citric acid cycle, requires MCM. Indeed, growth of *S. meliloti* on L-isoleucine and L-valine as the only carbon sources was dependent on the presence of the *bhbA* gene, which encodes MCM [91] (Figure S3).

We constructed an *S. meliloti* strain incapable of synthesizing cobalamin and lacking cobamide-dependent enzymes other than MCM to ensure that differential growth could be attributed solely to MCM selectivity for added cobamides (see Materials and Methods). We cultivated this strain with L-isoleucine and L-valine as sole carbon sources in medium supplemented with different cobamides in their cyanylated (CN) forms, which is the form typically used for *in vivo* growth assays. Under these growth conditions, the maximum growth yield (OD_600_) achieved at high concentrations of all of the cobamides was indistinguishable (Figure S4A-G). However, the concentration of cobamides required to achieve half of the maximal OD_600_ (EC_50_) differed based on the cobamide provided (Figure 5). Consistent with the binding data, CNCbl had the lowest EC_50_ value. EC_50_ values for CN[Bza]Cba and CN[2-MeAde]Cba were 5-fold higher than CNCbl, and other cobamides had EC_50_ values two orders of magnitude higher than CNCbl.

**Figure 5:**
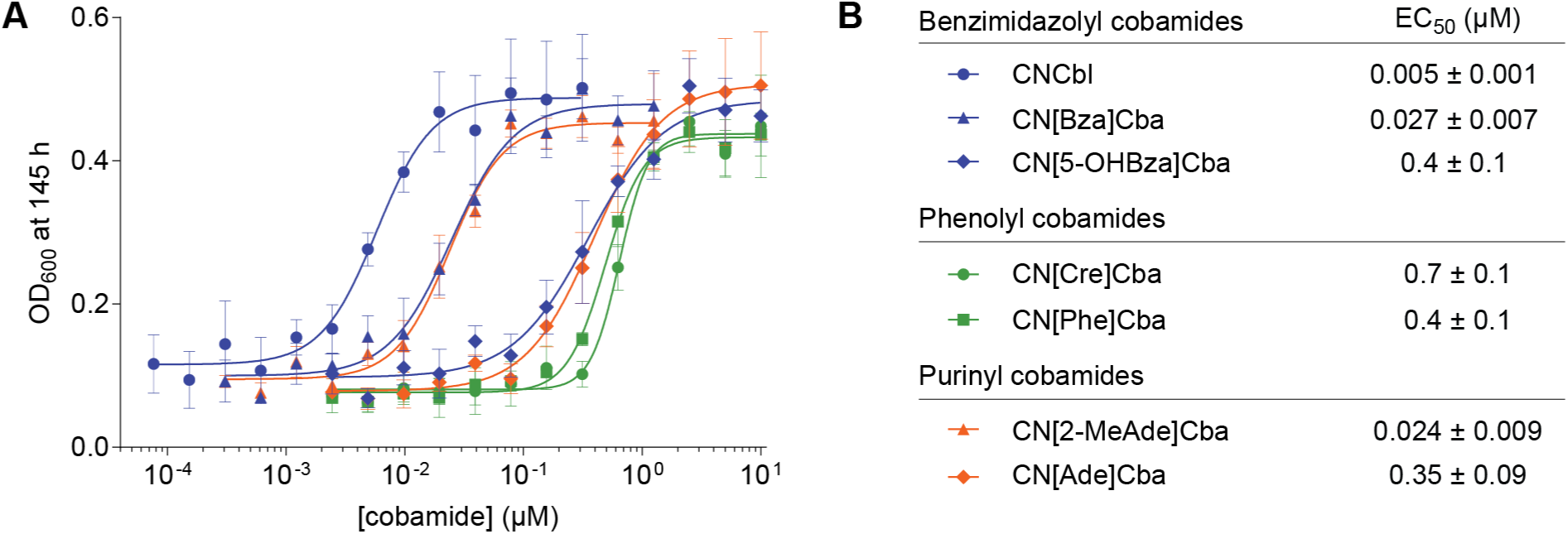
MCM-dependent growth of *S. meliloti cobD*::*gus* Gm^R^ *metH*::Tn*5* Δ*nrdJ* pMS03-*nrdAB*_*Ec*_^+^ with various cobamides (see also Figures S3, S4, S5). (A) Dose dependence of growth based on OD_600_ at 145 h. Data points and error bars represent the mean and standard deviation, respectively, of three biological replicates from a single experiment. EC_50_ values reported in (B) are the average and standard deviation of five or more biological replicates across two or more independent experiments.

With the notable exception of the phenolyl cobamides, differences in the EC_50_ values of cobamides *in vivo* qualitatively correlated with the binding selectivity that we observed *in vitro* (Figure 2). Among benzimidazolyl cobamides, EC_50_ values increased from cobalamin to [Bza]Cba to [5-OHBza]Cba, consistent with the relative binding affinities of these cobamides. Similarly, [2-MeAde]Cba, which had an intermediate binding affinity for *Sm*MCM, had a lower EC_50_ value than [Ade]Cba, which did not bind to *Sm*MCM at low micromolar concentrations *in vitro.* The ability of [5-OHBza]Cba and [Ade]Cba to support growth suggests that these cobamides can bind *Sm*MCM at concentrations higher than those tested *in vitro*; a control experiment with an *S. meliloti* strain lacking MCM rules out the possibility that high concentrations of cobamides (10 µM) abiotically enable growth on isoleucine and valine (Figure S3).

We considered the possibility that differences in cobamide internalization by *S. meliloti* could also influence the EC_50_ measurements shown in Figure 5. When *S. meliloti* cultures were supplemented with equimolar amounts of CNCbl, CN[Ade]Cba, or CN[Cre]Cba, the concentration of cobalamin extracted from the cellular fraction was 2- to 3-fold higher than [Ade]Cba and 5- to 6-fold higher than [Cre]Cba (Figure S5A). This result suggests that cobamides are differentially internalized or retained by the cells. However, MCM-dependent growth does not correlate with intracellular cobamide concentrations, as intracellular concentrations of cobalamin comparable to those of [Ade]Cba and [Cre]Cba supported *S. meliloti* growth to high densities (Figure S5). Therefore, the high EC_50_ of CN[Ade]Cba relative to CNCbl is more likely attributable to enzyme selectivity. Additional factors that could explain the high EC_50_ values of the phenolyl cobamides are considered in the Discussion.

### Identification of structural elements that interfere with cobamide binding

Given the apparent importance of MCM cobamide binding selectivity for the cobamide-dependent growth of *S. meliloti*, we pursued a more mechanistic understanding of how lower ligand structure affects cobamide binding. When cobamides are bound to MCM, the lower ligand is surrounded by protein residues [61, 97]. Therefore, the reduced affinity of certain cobamides for the enzyme could be a result of exclusion of their lower ligands from this binding pocket because of steric or electrostatic repulsion. We hypothesized that the poor binding of the purinyl cobamides Ado[Ade]Cba and Ado[2-MeAde]Cba is due to the presence of the exocyclic amine (N10) based on several observations: 1) Ado[5-OHBza]Cba, which also contains a polar functional group, had impaired binding to *Sm*MCM (Figure 2A, C). 2) In the crystal structure of of *Hs*MCM [97], residues Phe722 and Ala731, which are conserved in *Sm*MCM, would be expected to electrostatically occlude the exocyclic amine of [Ade]Cba (Figure S6A, asterisk). In contrast, polar residues Tyr and Ser occupy these positions in *Ec*MCM, which has higher affinity for purinyl cobamides (Figure S2A). 3) Structural modeling of Ado[Ade]Cba bound to *Hs*MCM, which shares 59% amino acid identity to *Sm*MCM in the B_12_-binding domain, suggests a significant displacement of the adenine lower ligand relative to the position of the lower ligand of AdoCbl, accompanied by significant expansion of the lower ligand binding pocket, which would be an unlikely conformation for the protein to adopt (Figure S6B-D).

To test the importance of the exocyclic amine of adenine in cofactor exclusion, we produced an unsubstituted purinyl cobamide, Ado[Pur]Cba [39]. Ado[Pur]Cba also had low affinity for *Sm*MCM (Figure 6A, B), suggesting that the exocyclic amine of adenine is not a major cause of binding exclusion. Consistent with this result, two novel benzimidazolyl cobamides, Ado[7-MeBza]Cba and Ado[7-AmBza]Cba, bound *Sm*MCM with comparable affinities to other benzimidazolyl cobamides (Figure 6A, B), despite being functionalized at the position analogous to N10 of adenine. Rather, these results suggest that the presence of nitrogens in the six-membered ring of the lower ligand interferes with binding. To test this hypothesis directly, we produced three novel cobamide analogs that contain at least one nitrogen in the six-membered ring of the lower ligand base. Comparison of the binding of Ado[6-MePur]Cba and Ado[7-MeBza]Cba (Figure 6A, B) supported a role of ring nitrogens in binding inhibition, and comparison of binding affinities between Ado[Bza]Cba and Ado[5-AzaBza]Cba (Figure 2A, C and Figure 6A, B, respectively), and between Ado[7-AmBza]Cba and Ado[3-DeazaAde]Cba (Figure 6A, B), revealed that a single nitrogen atom in the six-membered ring of the lower ligand was sufficient to severely impair binding.

**Figure 6:**
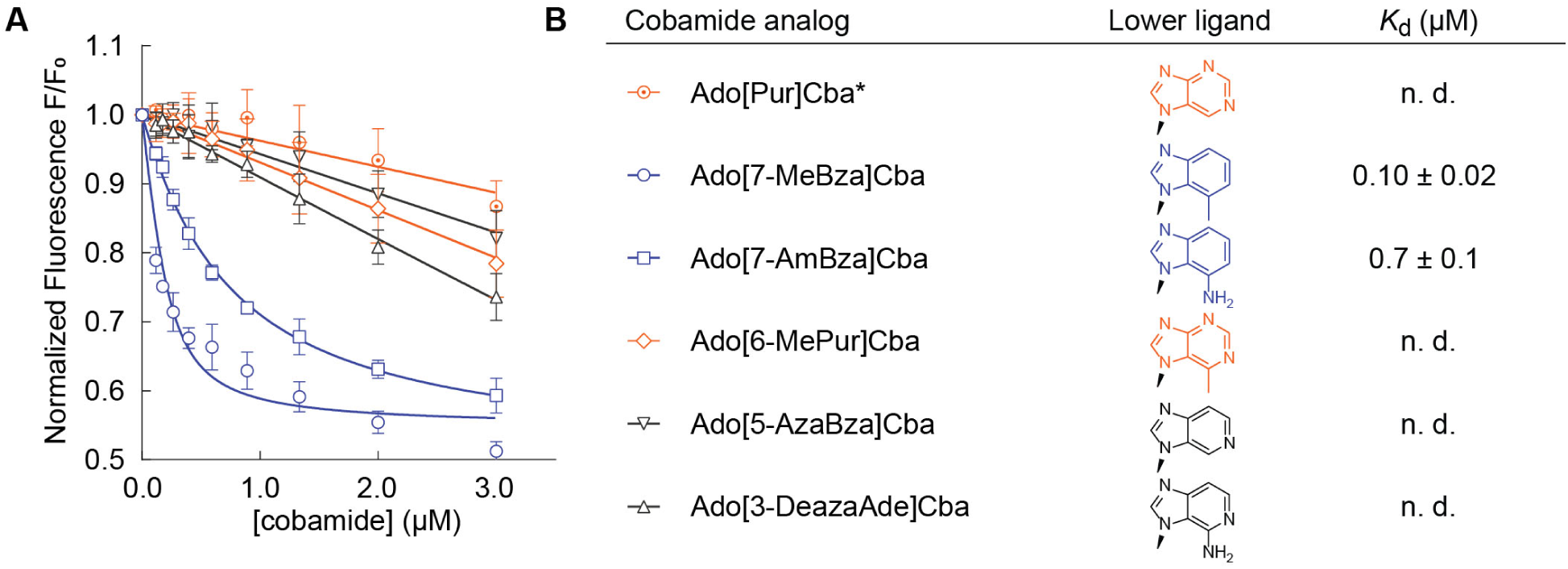
Binding of cobamide analogs to *Sm*MCM (see also Figure S7). (A) Fluorescence decrease of *Sm*MCM when reconstituted with benzimidazolyl (blue), purinyl (orange), and azabenzimidazolyl (black) cobamide analogs. Data points represent the mean and standard deviation of three technical replicates from a single experiment. (B) *K*_d_ values for different cobamides, reported as the average and standard deviation of three or more independent experiments, each consisting of technical triplicates. “n. d.,” not determined, indicates that binding was too weak to determine *K*_d_. *While it was unreported at the time of our study, [Pur]Cba was discovered to be the cobamide naturally produced by *Desulfitobacterium hafniense* [39].

As it was recently discovered to be a naturally occurring cobamide [39], we tested the MCM-dependent growth of *S. meliloti* with [Pur]Cba. [Pur]Cba had a high EC_50_ value of 0.6 ± 0.2 µM (Figure S4H), further supporting the correlation between binding and growth that we previously observed for benzimidazolyl and purinyl cobamides.

## Discussion

Cobamides are distinct from other cofactors in their extensive structural diversity, with over a dozen forms that differ in the lower ligand base and nucleotide loop. How cobamide lower ligand structure influences the activity of cobamide-dependent enzymes has not been extensively explored. Here, we report a systematic analysis of the effects of cobamide lower ligand structure on the function of a model cobamide-dependent enzyme, MCM. Our results show that MCM exhibits varied affinities for different cobamides, and that this selectivity is linked to the physiology of the organism.

Our results show that the major determinant of cobamide selectivity in *Sm*MCM is binding, with small changes in the lower ligand capable of dramatically altering the binding affinity of a cobamide. One explanation for these differences is that the chemical compatibility between the lower ligand base and the binding pocket of the protein strongly influences the binding affinity of cobamides; repulsion of the lower ligand on the basis of electrostatics could reduce the binding affinity of cobamides to MCM. While the structure of *Sm*MCM has not been determined, a model generated by sequence alignment to *Hs*MCM suggested a highly hydrophobic lower ligand binding pocket. Consistent with this, we observed higher affinity of cobamides with hydrophobic lower ligands to *Sm*MCM, as well as interference of ring nitrogens with cobamide binding.

On the other hand, sequence alignments suggested that many of the hydrophobic residues predicted to immediately surround the lower ligand are conserved between diverse MCM orthologs that differ in cobamide selectivity. Assuming that the arrangement of the lower ligand binding pocket is similar across MCM orthologs, this suggests that interactions within the lower ligand binding pocket are not sufficient to account for selectivity. In a similar vein, examination of the residues surrounding the lower ligand in the cobamide-bound structures of reductive dehalogenases does not reveal the basis of exclusion of certain cobamides [49]. These observations suggest that the lower ligand may have an unknown role in the binding of cobamides to MCM. Consistent with this idea, studies of the kinetics and pH dependence of AdoCbl binding to *P. shermanii* MCM suggest a pre-association step, wherein a cofactor-protein complex is formed prior to displacement of the lower ligand of the cofactor by a histidine residue in the protein [89]. The nature of this complex is unknown, but potential interactions between the lower ligand and this conformation of the enzyme could provide an opportunity for lower ligand structure to impact the outcome of binding.

Our analysis of MCM orthologs from *E. coli* and *V. parvula* demonstrates that variations in cobamide selectivity have evolved in organisms with different physiologies. The cobamide selectivity patterns in the three MCM orthologs we examined correlate with the physiologies of the bacteria in two ways. First, in all three cases, each MCM ortholog has highest affinity for the native cobamide produced by the organism, suggesting that cobamide biosynthesis and selectivity of cobamide-dependent enzymes have coevolved. Second, *Sm*MCM is more selective than *Ec*MCM and *Vp*MCM, and these differences in selectivity correlate with differences in cobamide biosynthesis, acquisition, and use in these organisms. *S. meliloti* synthesizes cobalamin *de novo* and is incapable of attaching purinyl and phenolyl lower ligands to cobamide precursors [96]. Thus, its cobamide-dependent enzymes have likely evolved to function best with cobalamin. In contrast, *E. coli* does not synthesize cobamides *de novo* and instead relies on the importer BtuBFCD to acquire cobamides from the environment [105, 106]. Alternatively, *E. coli* can produce a variety of benzimidazolyl and purinyl cobamides when provided with precursors [95], making the ability to use multiple cobamides likely advantageous. Like *S. meliloti, V. parvula* synthesizes cobamides *de novo*, but can produce both benzimidazolyl and phenolyl cobamides [96, 107] and also encodes membrane transport components adjacent to cobalamin riboswitches [108], which are likely to be cobamide importers [52, 109]. Thus, the ability of *Vp*MCM to bind diverse cobamides is similarly consistent with its physiology.

Relative to cobamide binding selectivity, our results suggest that effects of lower ligand structure on the catalytic activity of MCM are minor. Among the cobamides we tested, the maximum differences in *Sm*MCM turnover were 3-fold. We did not observe inhibition of MCM activity with any cobamides, in contrast to the strong inhibition that has been observed with analogs containing variations in the upper ligand or central metal, known as antivitamins [110-112].

In addition to elucidating the biochemical basis of cobamide selectivity in MCM, a major aim of our work was to link biochemical selectivity with cobamide-dependent growth. Our results with benzimidazolyl and purinyl cobamides support the hypothesis that enzyme selectivity is a major determinant of cobamide-dependent growth. Interestingly, although phenolyl cobamides bound *Sm*MCM with high affinity and supported catalysis *in vitro*, high concentrations were required to support growth of *S. meliloti*. This discrepancy can be partially explained by poorer internalization or retention of these cofactors as compared to cobalamin (Figure S5). The observation that the intracellular cobamide concentrations were 50- to 190-fold higher than the amount added to the growth medium (Figure S5A) suggests that cobamides could be internalized by an uptake mechanism that favors cobalamin, distinct from both BtuBFCD and ECF-CbrT [113, 114], both of which are absent from *S. meliloti*. Thus, we propose a model in which the cobamide-dependent growth of bacteria is influenced not only by binding selectivity of cobamide-dependent enzymes, but also by cobamide import (Figure 7). The lower effectiveness of phenolyl cobamides in supporting growth of *S. meliloti* could additionally be explained by inefficient adenosylation of these cobamides *in vivo*, as MCM requires the adenosyl upper axial ligand for activity. Whether or not adenosyltransferase enzymes, specifically CobA and PduO [115, 116] in *S. meliloti*, are selective with respect to lower ligand structure is unknown.

**Figure 7:**
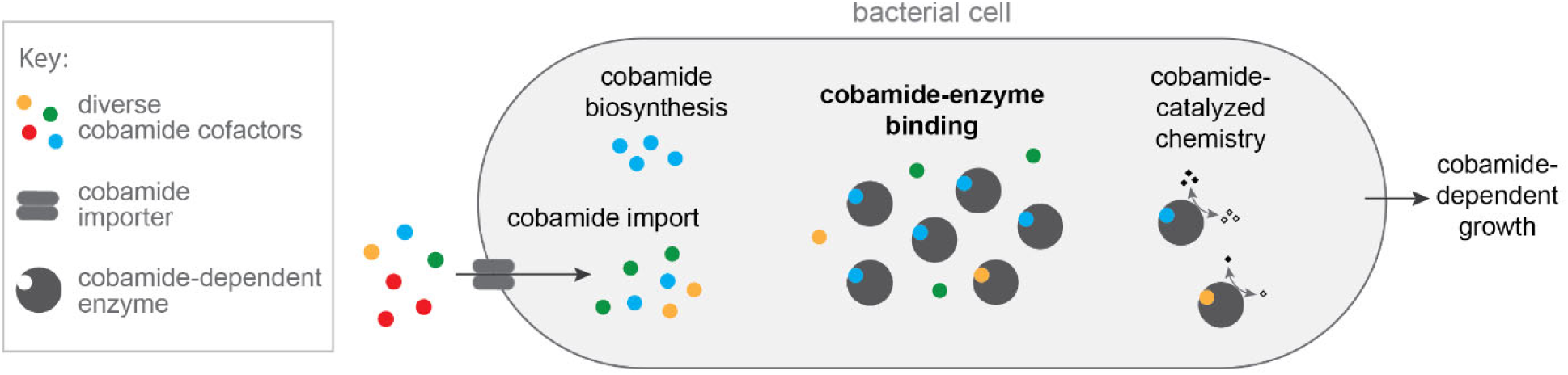
Model describing biochemical determinants of cobamide-dependent growth in bacteria. Cobamides differentially impact bacterial growth due to selective cobamide import and biosynthesis, cobamide-binding selectivity of cobamide-dependent enzymes, and cobamide-dependent catalysis. For MCM-dependent growth of *S. meliloti*, cobamide-binding selectivity is most strongly correlated with cobamide-dependent growth of the organism.

We and others have proposed the possibility of manipulating microbial communities using cobamides by taking advantage of the differential cobamide-dependent growth of bacteria [39, 117-119]. Cobamides are predicted to mediate microbial interactions that are critical to the assembly of complex communities [33, 41, 45, 50, 120-123], so the ability to selectively inhibit or promote the growth of particular species using corrinoids with various lower ligands could be applied to alter the composition of microbial communities in ways that could promote environmental and human health. This possibility hinges on the ability to predict which cobamides support or inhibit growth of an organism of interest, which requires an understanding of the major biochemical determinants of growth. We observed here that the cobamide binding selectivity of a model base-off cobamide-dependent enzyme correlates with growth to a large extent. Thus, uncovering protein residues that confer selectivity would enable prediction of selectivity in cobamide-dependent enzymes, thereby facilitating prediction of the cobamide requirements of organisms of interest. Furthermore, our results suggest that additional steps of cobamide trafficking may be important determinants of cobamide-dependent growth. Future studies to understand how these various steps depend on cobamide structure will ultimately allow us to better understand, predict, and manipulate microbial interactions.

## Materials and Methods

### Chemical reagents

Chemicals were obtained from the following sources: 5’-chloro-5’-deoxyadenosine, Santa Cruz Biotechnology; 4-methylbenzimidazole, Accela; 5-methyl-1H-benzimidazole, ACROS Organics; phenol, J. T. Baker; zinc metal, Fisher Scientific; 5-methoxybenzimidazole, purine, and *para*-cresol, Alfa Aesar; methylmalonyl-CoA, methylmalonic acid, coenzyme A, adenosylcobalamin (coenzyme B_12_), cyanocobalamin, dicyanocobinamide, 6-methylpurine, 1H-imidazo[4,5-c]pyridine-4-amine (3-deazaadenine), benzimidazole, adenine hemisulfate, 5- azabenzimidazole, 1H-benzo[d]imidazol-7-amine (4-aminobenzimidazole), 2-methyl-1H-purine-6-amine (2-methyladenine), and bovine serum albumin (BSA), Sigma.

### Molecular cloning, protein expression and purification

*Sm*MCM (locus SM_b20757, *bhbA*) was expressed from the pET28a vector, with an N-terminal hexahistidine (6xHis) tag, in *E. coli* BL21(DE3)pLysS (cloning primers listed in Table S1). The expression strain was grown to an optical density at 600 nm (OD_600_) of 0.6-0.8 at 37 °C, cooled on ice for 15 min, and induced with 1 mM IPTG for 2.5 h at 37 °C. Cells were lysed by sonication in 25 mM Tris-HCl pH 8.0, 300 mM NaCl, 10 mM imidazole, with 0.5 mM PMSF, 1 µg/mL leupeptin, 1 µg/mL pepstatin, and 1 mg/mL lysozyme. Clarified lysate was treated with 0.05% polyethyleneimine. An ÄKTA Pure 25 Fast Protein Liquid Chromatography (FPLC) system was used to purify the protein over a GE 5 mL HisTrap HF column, using a gradient of 21 to 230 mM imidazole in the lysis buffer. Purified protein was dialyzed into 25 mM Tris-HCl pH 8.0, 300 mM NaCl, 10% glycerol and concentrated with a Vivaspin 10,000 MWCO protein concentrator. Final protein concentration was determined by A_280_ using the theoretical extinction coefficient 55810 M^−1^ cm^−1^ [124]. *Ec*MCM (locus b2917, *scpA*, previously *sbmA*) was expressed with an N-terminal 6xHis tag from a pET28a vector in *E. coli* BL21(DE3), by induction at OD_600_ 0.6-0.8 with 0.1 mM IPTG, for 3.5 h at 30 °C. The protein was purified as described above and the final concentration was determined by Coomassie-stained SDS-PAGE, using BSA as a standard.

The *V. parvula* genome has two MCM annotations: a heterotetramer (loci Vpar_RS06295, Vpar_RS06290) and a heterodimer (loci Vpar_RS09005, Vpar_RS09000). The functionality of both homologs was tested by complementation in *S. meliloti*. The two putative *Vp*MCM enzymes were cloned into the pTH1227 vector and transferred by conjugation into an *S. meliloti bhbA*::Tn*5* mutant. Complementation was assessed by growth in M9 liquid medium containing L-isoleucine and L-valine (see “*S. meliloti* growth assays” for additional details). *S. meliloti* co-expressing Vpar_RS09005 and Vpar_RS09000 showed identical growth to a strain expressing *Sm*MCM from pTH1227 and was selected for *in vitro* studies.

The α subunit of *Vp*MCM (encoded by Vpar_RS09005) was expressed with an N-terminal 6xHis tag from the pET-Duet expression vector in *E. coli* BL21(DE3). Protein expression was induced with 520 µM IPTG for 6 h at 30 °C. The protein was batch purified by nickel affinity and subsequently purified by FPLC using a HiTrapQ column with an NaCl gradient from 50 to 500 mM in 20 Tris-HCl pH 8.0, 10% glycerol. The β subunit of *Vp*MCM (encoded by Vpar_RS09000) was expressed separately with an N-terminal 6xHis tag from the pET-Duet expression vector in *E. coli* BL21(DE3). Expression was induced with 1 mM IPTG for 22 h at 16 °C and the protein was purified using nickel-affinity chromatography as described for *Sm*MCM. Purified protein was dialyzed into 25 mM Tris-HCl pH 8.0, 300 mM NaCl, 10% glycerol, and 1 mM β-mercaptoethanol. Concentration of α and β subunits was determined by absorbance at 280 nm (A_280_) using the theoretically calculated extinction coefficients 75290 M^−1^ cm^−1^ and 74260 M^−1^ cm^−1^, respectively [124]. Equimolar amounts of α and β subunits were combined during the setup of fluorescence binding assays.

*E. coli* thiokinase containing an N-terminal 6xHis tag was expressed from a vector provided by Gregory Campanello from the laboratory of Ruma Banerjee. Expression was induced with 1 mM IPTG in *E. coli* BL21(DE3)pLysS at 28 °C for 3 h. The protein was purified as a heterodimer using nickel-affinity chromatography as described above. His-tagged *Rhodopseudomonas palustris* MatB [125] was expressed from a pET16b expression plasmid provided by Omer Ad from the laboratory of Michelle Chang. The protein was overexpressed in *E. coli* BL21(DE3) at 16 °C overnight, after induction with 1 mM IPTG, and purified by nickel affinity chromatography as indicated above. Thiokinase and MatB concentrations were determined by Coomassie-stained SDS-PAGE, using BSA as a standard.

### Guided biosynthesis, extraction, and purification of cobamides

*Sporomusa ovata* strain DSM 2662 was used for the production of its native cobamide, [Cre]Cba, and for production of [Phe]Cba, [5-MeBza]Cba, [Bza]Cba, [5-OHBza]Cba, [7-MeBza]Cba, and [7-AmBza]Cba, by guided biosynthesis as previously described [44]. 5-OHBza was synthesized as described [96]. *Salmonella enterica* serovar Typhimurium strain LT2 and *Propionibacterium acidipropionici* strain DSM 20273 were used for production of [Ade]Cba [47, 126]. [2-MeAde]Cba, [Pur]Cba, [5-AzaBza]Cba, [3-DeazaAde]Cba, and [6-MePur]Cba were produced by guided biosynthesis in *P. acidipropionici*. Cobamides were extracted as previously described [47] and purified by High Performance Liquid Chromatography (HPLC) using previously published methods [47, 96, 127] as well as additional methods listed in Table S2. In many cases more than one method was required to achieve high purity. Identity of cobamides was confirmed by Liquid Chromatography (LC) coupled to Mass Spectrometry (MS) using an Agilent 1260 LC/6120 quadrupole MS instrument. Lower ligand orientation in the novel cobamides [7-MeBza]Cba, [7-AmBza]Cba, [3-DeazaAde]Cba, and [6-MePur]Cba was inferred based on their absorbance spectra, which reveal a base-on conformation in the cyanylated form (Figure S7). The orientation of the lower ligands in [Pur]Cba and [5-AzaBza]Cba was not determined.

### Chemical adenosylation of cobamides

Cobamide adenosylation was performed as previously described [127, 128]. Briefly, cobamides at concentrations 0.5 – 1 mM were reduced with activated zinc metal under anaerobic conditions, with vigorous stirring for 0.5 – 2 h. 5’-chloro-5’-deoxyadenosine was added and adenosylation was allowed to proceed for 1 – 3 h in the dark. The progress of the reaction was monitored by HPLC. Following adenosylation, cobamides were desalted using a C18 SepPak (Waters), purified by HPLC, desalted again, dried, and stored at −20 °C or −80 °C.

### Cobamide quantification

Purified cobamides were dissolved in water and quantified by UV-Vis spectrophotometry on a BioTek Synergy 2 plate reader using the following extinction coefficients: for cyanylated benzimidazolyl cobamides, ε_518_ = 7.4 × 10^3^ M^−1^ cm^−1^ [129]; for cyanylated purinyl cobamides, ε_548_ = 7.94 × 10^3^ M^−1^ cm^−1^ [130]; for cyanylated phenolyl cobamides, ε_495_ = 9.523 × 10^3^ M^−1^ cm^−1^ [131]; for adenosylated benzimidazolyl cobamides (AdoCbl, Ado[5-MeBza]Cba, Ado[Bza]Cba, Ado[5-OHBza]Cba, Ado[7-MeBza]Cba, and Ado[7-AmBza]Cba), which are predominantly base-on in water, ε_522_ = 8.0 mM^−1^ cm^−1^ [129]; for adenosylated purinyl and phenolyl cobamides (Ado[Ade]Cba, Ado[2-MeAde]Cba, Ado[Pur]Cba, Ado[Cre]Cba, and Ado[Phe]Cba), which are predominantly base-off in water, ε_458_ = 8.8 mM^−1^ cm^−1^ [130]; for adenosylated azabenzimidazolyl cobamides (Ado[3-DeazaAde]Cba, Ado[5-AzaBza]Cba and Ado[6-MePur]Cba), which are a mixture of base-on and base-off in water, the concentration was estimated from the average of concentrations calculated using the extinction coefficients above.

### Fluorescence Binding Assays

An *in vitro* assay previously described for measuring binding of AdoCbl to *P. shermanii* MCM [89] was adapted to a 96-well format: MCM (0.2 µM) was combined with a range of cobamide concentrations (as specified in each experiment) in a black 96-well plate in 50 mM potassium phosphate pH 7.5 with 1 mM DTT, on ice. All steps involving cobamides were conducted in the dark. The plate was centrifuged for 1 min at 3800 rpm to level the surface of the liquid in each well. The plate was then incubated for 40 min at 30 °C to allow binding, with a brief shaking step after 30 min. Preliminary experiments showed that this time is sufficient for equilibration. Following incubation, fluorescence emission at 340 nm (5 nm slit width) was measured upon excitation at 282 nm (5 nm slit width) using a Tecan Infinite M1000 PRO Plate Reader. Fluorescence, normalized to the initial value, was plotted as a function of cobamide concentration, and fit to the following equation [132]:

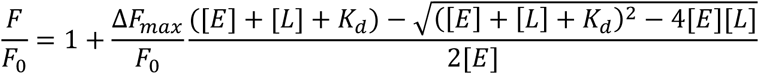

where *F* is fluorescence, *F*_0_ is initial fluorescence, [*E*] is total enzyme concentration, [*L*] is total ligand concentration, and *K*_*d*_ is the binding dissociation constant.

### Filtration binding assay

Cobamides (10 µM) with and without MCM (15 µM) were incubated in 50 mM Tris-phosphate buffer pH 7.5 at 30 °C for 40 min, transferred to Nanosep 10K Omega centrifugal devices (Pall Corporation), and centrifuged for 5 minutes at 13,900 × g to separate unbound cobamides from enzyme-bound cobamides. The UV-Vis spectra of the filtrates were recorded on a BioTek Synergy 2 plate reader.

### Structural modeling

A model of *Sm*MCM was generated using the Swiss-Model software [133] based on the known crystal structure of *Homo sapiens* MCM (*Hs*MCM) (PBD ID: 2XIJ) [97]. No major differences were observed in the B_12_-binding domain between *Sm*MCM models generated from *Hs*MCM and *Propionibacterium freudenrichii* MCM (PDB ID: 4REQ) [61].

Maestro [134] was used to generate a model of *Hs*MCM bound to [Ade]Cba. The initial structure of *Hs*MCM bound to cobalamin (PDB ID: 2XIQ) [97] was prepared using standard methods. A constrained energy minimization (atoms within 10 Å of cobalamin freely moving; atoms within a second 10 Å shell constrained by a force constant of 200; remaining structure frozen) was performed using MacroModel [135]. The structure of the lower ligand was then modified to adenine, and the constrained energy minimization was repeated to generate a model of the lower ligand binding pocket bound to [Ade]Cba.

### Enzymatic synthesis of (R)-methylmalonyl-CoA

(*R*)-methylmalonyl-CoA synthesis reactions contained the following in 10 mL: 100 mM sodium phosphate pH 7.5, 20 mM MgCl_2_, 5 mM ATP, 10 mM methylmalonic acid, 2 mM coenzyme A, 5 mM β-mercaptoethanol, and 1.5 µM purified MatB protein. After combining ingredients on ice, the reaction was incubated at 37 °C for 1 h. The reaction was then frozen in liquid nitrogen and lyophilized. To purify (*R*)-methylmalonyl-CoA, the dried reaction mixture was resuspended in 3.2 mL water and the protein was precipitated with 200 µL trichloroacetic acid; precipitate was pelleted; supernatant was neutralized with 200 µL of 10 M NaOH; and salts and remaining starting materials were removed using a C18 SepPak column (Waters) (loaded in 0.1% formic acid, washed with water, methylmalonyl-CoA eluted with 50% methanol in water). Formation of (*R*)-methylmalonyl-CoA was initially verified by ^1^H NMR and in subsequent preparations by HPLC (Table S3). The concentration of (*R*)-methylmalonyl-CoA was determined using an extinction coefficient of 12.2 mM^−1^ cm^−1^ at 259 nm.

### MCM activity assays

A thiokinase-coupled, spectrophotometric MCM activity assay was adapted from previous work [136], except that ADP was used instead of GDP, and the experiment was conducted in 96-well plates. Final concentrations of reagents in the assays are as follows: Tris-phosphate buffer pH 7.5, 50 mM; DTNB, 400 µM; ADP, 1 mM; MgCl_2_, 10 mM; (*R*)-methylmalonyl-CoA, 0 – 4 mM; thiokinase, 5 µM; MCM, 50 nM; and cobamides, 2 µM. Preliminary experiments were conducted to ensure that concentrations of thiokinase, DTNB, and cobamides were not rate limiting.

Three separate mixes were prepared, all in 1X Tris-phosphate buffer: an assay mix containing DTNB, ADP, and MgCl_2_, a substrate mix containing (*R*)-methylmalonyl-CoA, and an enzyme mix containing thiokinase, MCM, and cobamides. All steps involving cobamides were conducted in the dark. The assay and enzyme mixes were prepared as a master mix and aliquoted into 96-well plates; substrate mixes were prepared in individual wells, in triplicate. All components were incubated at 30 °C for 40 minutes to equilibrate temperature and allow pre-binding of cobamides and MCM. After incubation, one replicate at a time, the substrate mix was added to the assay mix, followed by the enzyme mix. Absorbance at 412 nm (A_412_) was recorded immediately after addition of enzyme and for 1-3 minutes, every 3 seconds, on a BioTek Synergy 2 plate reader. The increase in A_412_ in reactions lacking substrate was subtracted from all readings, to account for reactivity of DTNB with thiols on protein surfaces. A_412_ values were converted to concentration of free CoA using a pathlength correction determined for the reaction volume and extinction coefficient of 14150 M^−1^ cm^−1^.

### S. meliloti growth assays

MCM-dependent growth experiments were performed with *S. meliloti* strain Rm1021 *cobD*::*gus* Gm^R^ *metH*::Tn*5* Δ*nrdJ* pMS03-*nrdAB*_*Ec*_^*+*^, which lacks cobamide-dependent enzymes other than MCM and does not synthesize cobalamin. *cobD* is required for cobalamin biosynthesis [137], *metH* encodes methionine synthase [137-139], and *nrdJ* encodes ribonucleotide reductase [140]. Because *nrdJ* is essential, the *E. coli* cobamide-independent ribonucleotide reductase encoded by *nrdA* and *nrdB* was expressed from the pMS03 plasmid [141]. The strain was pre-cultured in M9 medium [142] (modified concentration of MgSO_4_: 1 mM) containing 0.1% sucrose, 2 g/L isoleucine, 2 g/L valine, 1 g/L methionine, and 20 µg/mL gentamycin, shaking at 30 °C. After two days, cells were washed and diluted to an OD_600_ of 0.02 into M9 medium containing 4 g/L isoleucine, 4 g/L valine, 1 g/L methionine, 20 µg/mL gentamycin, and cobamides at various concentrations as indicated for each experiment, in 384-well plates. The plates were incubated at 30 °C for 145 h in a Biotek Synergy 2 plate reader with linear shaking at 1140 cpm. OD_600_ was measured in 1 h increments.

For quantification of intracellular cobamides in *S. meliloti*, the strain above was pre-cultured as described, diluted into 50 mL of M9 medium containing 0.2% sucrose and various cobamides, and grown for 48 h (until OD_600_ 0.6-0.8). Cobamides were extracted from cell pellets as previously described [47], using 5 mL of methanol containing 500 µg of potassium cyanide, and including a partial purification by means of a wash step with 20% methanol in water during the SepPak desalting procedure. Extracted cobamides were quantified by HPLC using peak areas at 525 nm and external standard curves, and cellular cobamide concentrations were calculated assuming 8 × 10^8^ cells/mL at OD_600_ 1.0 and cellular volume of 1 µm^3^.

## Acknowledgements

We thank current and past members of the Taga lab and Ruma Banerjee, Judith Klinman, Susan Marqusee, and David Savage for helpful discussions; Amrita Hazra, Amanda Shelton, Alexa Nicolas, Zachary Hallberg, Kristopher Kennedy, Joseph Maa, and Judith Klinman for critical reading of the manuscript; Anna Beatrice Grimaldo for help with guided biosynthesis of cobamide analogs; Amrita Hazra and Florian Widner for providing 5-OHBza; Victoria Innocent for assisting with cobamide accumulation assays; Ruma Banerjee, Gregory Campanello, Michelle Chang, and Omer Ad for providing expression plasmids; Kathleen Durkin for help with molecular modeling; and Krishna Niyogi and Arash Komeili for use of their equipment. This work was supported by NIH grants R01GM114535 and DP2AI117984 to MET. OS was also supported by the NIH Chemical Biology Training Grant T32 GM066698, JT by the UCSF work study program, and modeling work at the UC Berkeley Molecular Graphics and Computation Facility by NIH S10OD023532.

