## Supplementary Figures and Tables for "Cofactor selectivity in methylmalonyl-CoA mutase, a model cobamide-dependent enzyme"

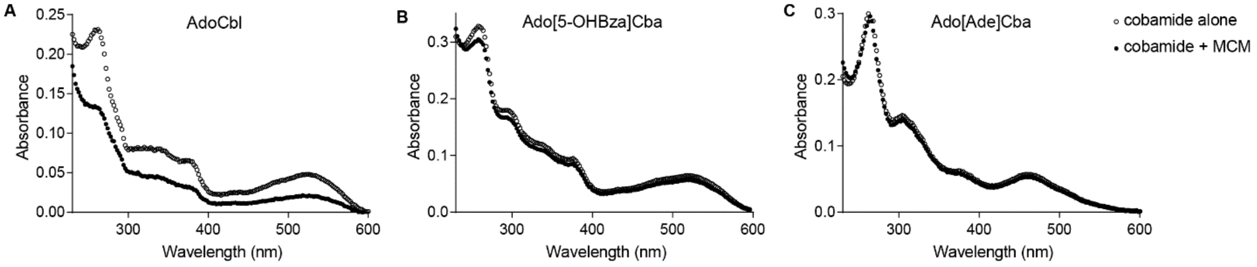


Figure S1: Filtration-based MCM binding assay. UV-Vis spectra of filtrate after pre-incubation of 10 μM (A) AdoCbl, (B) Ado[5-OHBza]Cba, and (C) Ado[Ade]Cba with and without *Sm*MCM (15 μM).

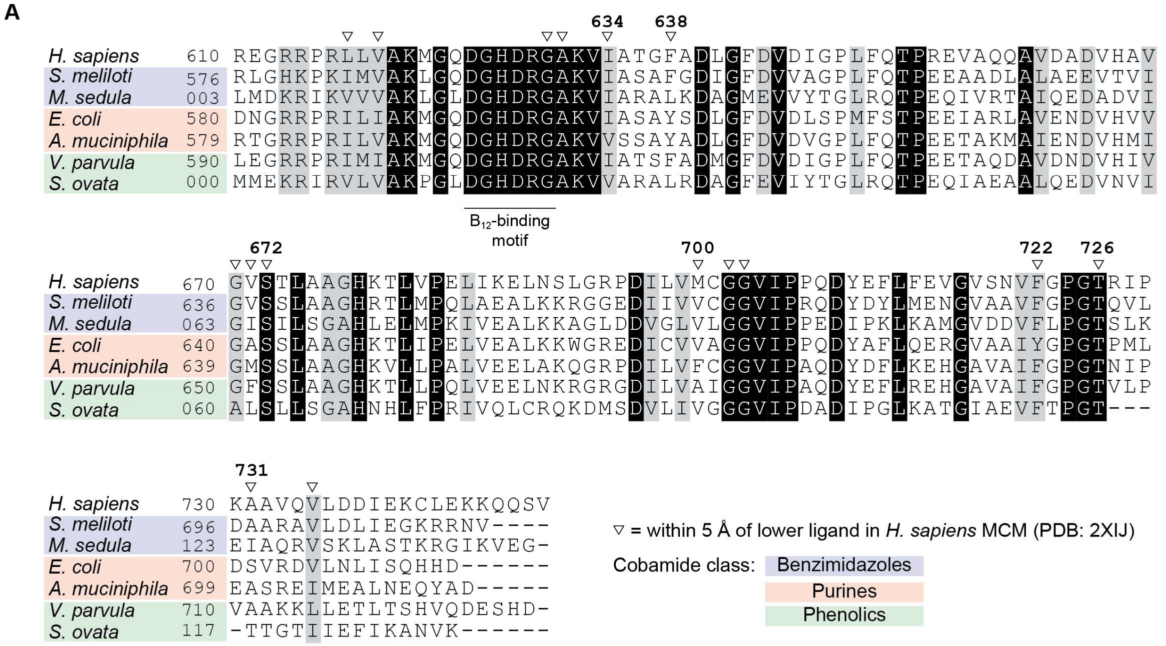


**B**

| % Identity | *H. sapiens* | *S. meliloti* | *M. sedula* | *E. coli* | *A. muciniphila* | *V. parvula* | *S. ovata* |
|---|---|---|---|---|---|---|---|
| *H. sapiens* | -- | 58 | 37 | 60 | 62 | 66 | 38 |
| *S. meliloti* | 58 | -- | 47 | 62 | 57 | 57 | 40 |
| *M. sedula* | 37 | 47 | -- | 43 | 39 | 37 | 57 |
| *E. coli* | 60 | 62 | 43 | -- | 62 | 63 | 39 |
| *A. muciniphila* | 62 | 57 | 39 | 62 | -- | 70 | 42 |
| *V. parvula* | 66 | 57 | 37 | 63 | 70 | -- | 38 |
| *S. ovata* | 38 | 40 | 57 | 39 | 42 | 38 | -- |

| Structural composition | |
|---|---|
| *H. sapiens* | $\alpha_2$ |
| *S. meliloti* | $\alpha_2$ |
| *M. sedula* | $\alpha_2\beta_2$ |
| *E. coli* | $\alpha_2$ |
| *A. muciniphila* | $\alpha\beta$ |
| *V. parvula* | $\alpha\beta$ |
| *S. ovata* | $\alpha_2\beta_2$ |

Figure S2: Sequence comparison of the $B_{12}$-binding domains of MCM orthologs. (A) Sequence alignment of the $B_{12}$-binding domains of MCM orthologs, generated using the MUSCLE alignment tool from EMBL-

EBI. Black and gray shading indicate amino acid identity and similarity, respectively. Sequences are colored by “cobamide class” based on cobamides biosynthesized by the organism or predicted cobamide use [1-5]. Residues numbered above the sequence alignment correspond to residues indicated in Figure S2. Locus tags of aligned proteins: *Homo sapiens* AAA59569, *Sinorhizobium meliloti* AAD13665, *Metallosphaera sedula* ABP96195, *Escherichia coli* WP_101348647, *Akkermansia muciniphila* WP_031931429, *Veillonella parvula* WP_004694550, *Sporomusa ovata* WP_021167215. (B) Percent identity matrix of the $B_{12}$-binding domains aligned in (A), as well as the structural composition of each MCM ortholog: $\alpha_2$, homodimer; $\alpha\beta$, heterodimer; $\alpha_2\beta_2$, heterotetramer.

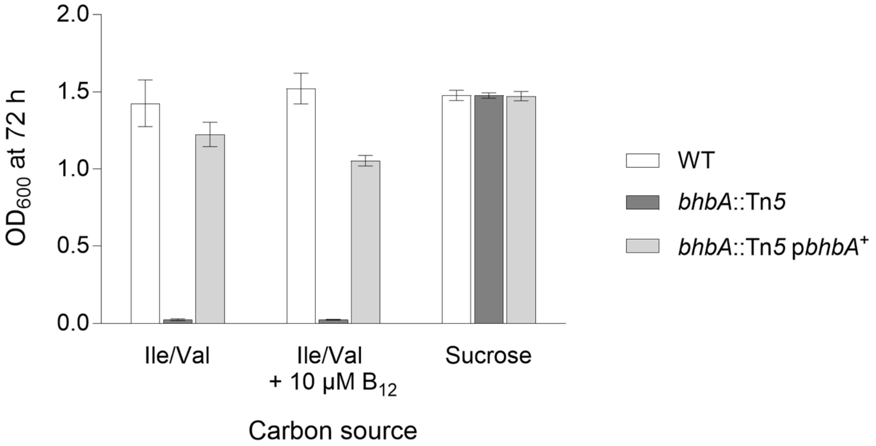


Figure S3: MCM-dependent growth of *S. meliloti*. Final density ($OD_{600}$), after 72 h of growth in M9 minimal medium, with 4 g/L L-isoleucine and 4 g/L L-valine (Ile/Val) or 2 g/L sucrose. *Sm*MCM is the gene product of the *bhbA* gene. p*bhbA*$^{+}$, complementation of the *bhbA*::Tn*5* mutation with the *S. meliloti bhbA* gene expressed in the pTH1227 vector [6]. Plot shows the mean and standard deviation of three biological replicates from a single experiment.

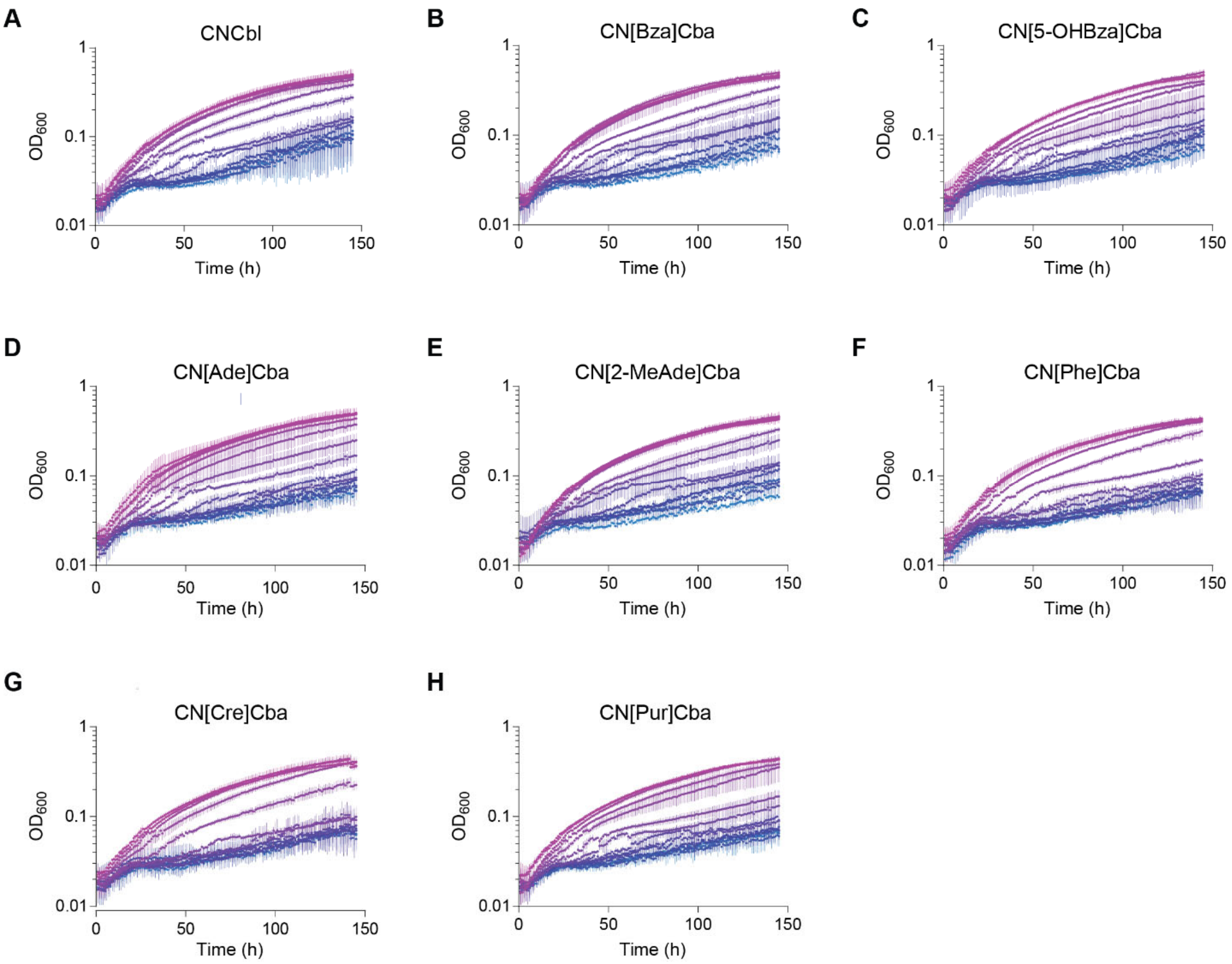


Figure S4: MCM-dependent growth of *S. meliloti cobD*::*gus* $Gm^R$ *metH*::Tn*5* Δ*nrdJ* pMS03-$nrdAB_{Ec}^+$ with different cobamides. Concentration decreases, by 2-fold dilutions, are shown from pink to blue. Plots show the mean and standard deviation of three biological replicates. Maximum concentrations tested (pink curves) are (A) CNCbl, 312.5 nM; (B) CN[Bza]Cba, 1.25 µM; (C) CN[5-OHBza]Cba, 10 µM; (D) CN[Ade]Cba, 10 µM; (E) CN[2-MeAde]Cba, 1.25 µM; (F) CN[Phe]Cba, 10 µM; (G) CN[Cre]Cba, 10 µM; and (H) CN[Pur]Cba, 10 µM.

A

| Cobamide | Concentration added to medium | Concentration in cells |
|---|---|---|
| CNCbl | 0.2 µM | 25 - 38 µM |
| | 0.1 µM | 5 - 17 µM |
| | 0.04 µM | 1 - 7 µM |
| CN[Ade]Cba | 0.2 µM | 11 - 14 µM |
| CN[Cre]Cba | 0.2 µM | 7 - 8 µM |

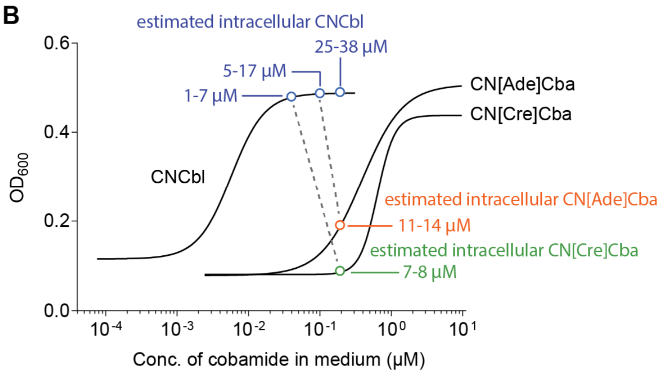


Figure S5: Quantification of cobamides internalized by *S. meliloti cobD*::*gus* Gm$^{R}$ *metH*::Tn*5* Δ*nrdJ* pMS03-*nrdAB$_{Ec}$*$^{+}$. (A) Cellular cobamide concentrations following 48 h of growth in M9 sucrose with the indicated concentrations of cobamides added to the medium. The range of concentrations measured in cell pellets was determined by HPLC analysis of corrinoid extractions from two or more independent experiments, each including biological duplicates. (B) A graphic illustrating the concentrations of different cobamides at which intracellular cobamide concentrations are comparable. Internalization data (colored) are overlaid onto dose-response curves from Figure 5 (black). Five points indicate cobamide concentrations at which cobamides were extracted and quantified; dotted lines connect conditions in which intracellular concentrations of different cobamides are approximately equal.

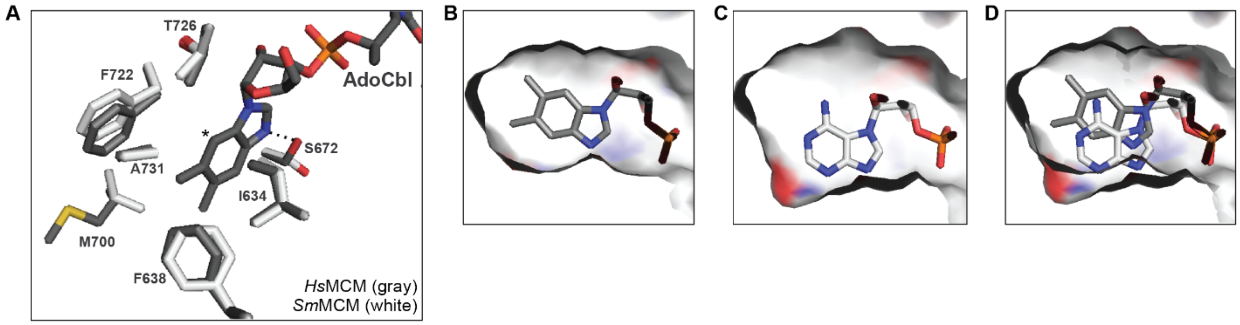


Figure S6: The lower ligand binding pocket of MCM. (A) Residues surrounding the lower ligand of AdoCbl in the X-ray crystal structure of *Homo sapiens* MCM (*Hs*MCM) (PDB: 2XIQ, gray). A model of *Sm*MCM, generated by sequence alignment and threading using Swiss-Prot, is overlaid in white. The asterisk marks the expected position of the exocyclic amine of [Ade]Cba. (B) Surface depiction of the lower ligand binding pocket of *Hs*MCM bound to cobalamin, after performing a constrained energy minimization. As expected, no major differences were observed between the energy minimized model and the original structure. (C) Surface depiction of the lower ligand binding pocket of *Hs*MCM modeled with [Ade]Cba bound, generated by changing the structure of the lower ligand in (B) and performing a constrained energy minimization. (D) Overlay of the structural models in B and C.

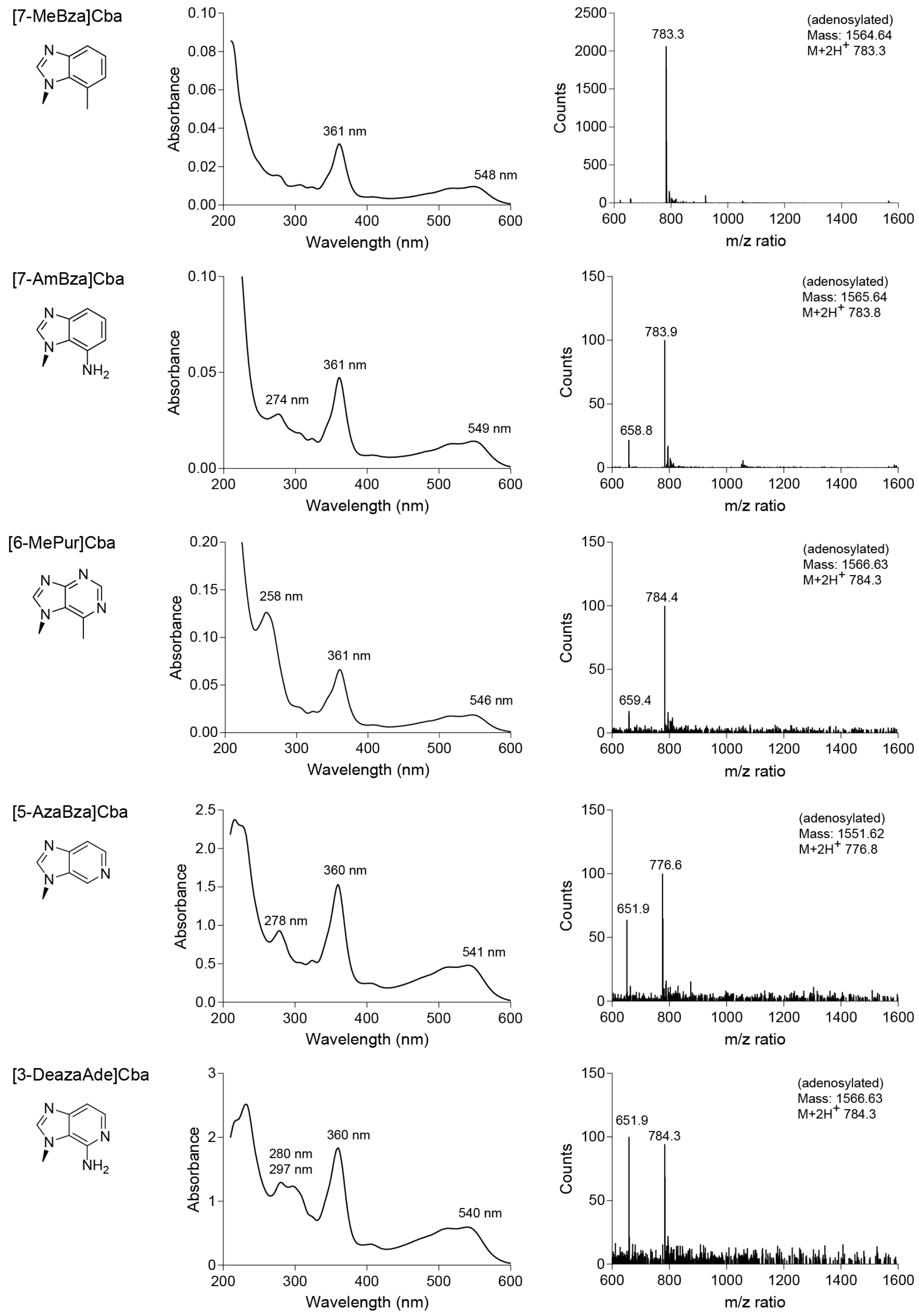


Figure S7: Spectral analysis of novel cobamide analogs. Left: Absorbance spectra of cyanylated cobamides, recorded during HPLC analysis of crude extracts. Labels indicate local absorbance maxima. Right: Mass spectra from HPLC-MS analysis of cobamides following adenosylation. Cobamides containing impurities were further purified by HPLC.

Table S1: Cloning primers.

| Construct | Primer sequence |
|---|---|
| pET28a-*bhbA*<br>(*Sm*MCM) | ccgcgcggcagccatatggctagcACCGAAAAGACCATCAAGGACTG<br>aagcttgtcgacggagctcgaattcTTACACGTTTCGCCGCTTGC |
| pET28a-*scpA*<br>(*Ec*MCM) | cgcgcggcagccatatgATGTCTAACGTGCAGGAGTGG<br>gtgcggccgcaagcttTTAATCATGATGCTGGCTTATCAG |
| pTH1227-Vpar_RS06295-Vpar_RS06290 | TTCACCTCGAGATCTATCGATGCATACCTGTTTTTTTTAGGAGGATGATGAAAAC<br>GCTTGAATTCGAGCTCCCGGGTACCTTATTTTACGTTTTCTTTAATAAAGTTAACGATATCGC |
| pTH1227-Vpar_RS09005-Vpar_RS09000 | TTCACCTCGAGATCTATCGATGCATGGTTTCGAGTTGAAAAGGAGGCA<br>GCTTGAATTCGAGCTCCCGGGTACCTCAGTCATGAGACTCGTCTTGAAC |
| pETDuet-Vpar_RS09005<br>(*Vp*MCM α) | cattggatcctATGTCTGACAAAAAGAC<br>ctaactcgagTCAGTCATGAGACTCGTC |
| pETDuet- Vpar_RS09000<br>(*Vp*MCM β) | cattggatcctATGTTTAAAAATC<br>ctaactcgagTCAGTCATGAG |

Table S2: HPLC methods for purification of cobamides.

| Compounds purified | Mobile Phases | Flow (mL/min) | Temp (˚C) | Gradient |
|---|---|---|---|---|
| Phenolyl, non-polar benzimidazolyl, and adenosylated cobamides | A: 0.1% formic acid in water<br>B: 0.1% formic acid in methanol | 2 | 30 | 25% B, 2 min<br>25 – 60% B, 24 min |
| Purinyl cobamides and azabenzimidazolyl cobamides | A: 0.1% formic acid in water<br>B: 0.1% formic acid in methanol | 1.5 | 15 | 10 – 30% B, 3 min<br>30% B, 13.9 min<br>30 – 37% B, 2.1 min |
| Purinyl cobamides and azabenzimidazolyl cobamides | A: 0.1% formic acid in water<br>B: 0.1% formic acid in methanol | 2 | 30 | 10 – 42% B, 20 min |
| [5-OHBza]Cba | A: 0.1% formic acid in water<br>B: 0.1% formic acid in methanol | 2 | 25 | 18 – 25% B, 2.5 min<br>25%, 22.5 min |
| β-adenosylcobinamide [7] | A: 10 mM sodium phosphate pH 7<br>B: acetonitrile | 2 | 25 | 2 – 23% B, 40 min |

Column: Zorbax Eclipse Plus C18, 9.4 x 250 mm, 5μm

Table S3: HPLC methods for analysis and purification of methylmalonyl-CoA

| Column | Mobile Phases | Flow (mL/min) | Temp (˚C) | Gradient |
|---|---|---|---|---|
| Eclipse XBD-C18<br>4.6 x 150 mm<br>5μm | A: 100 mM acetic acid, 100 mM sodium phosphate, pH 4.6<br>B: 100 mM acetic acid, 100 mM sodium phosphate, pH 4.6 18% methanol | 0.75 | 40 | 44% B, 14 min |
| Zorbax Eclipse Plus C18<br>9.4 x 250 mm<br>5μm | A: 0.1% formic acid in water<br>B: acetonitrile | 3 | 25 | 0 – 10% B, 30 min<br>10 – 70% B, 3 min |
